# Urban PM_2.5_ Impairs Blood–Brain Barrier Integrity and Enhances LOX-1 Expression in Human Brain Endothelial Cells

**DOI:** 10.64898/2026.01.29.702473

**Authors:** Elisabet Andersson, Trevor Wendt, Fanny Bergman, Christina Isaxon, Saema Ansar

## Abstract

**Introduction:** Ambient air pollution, especially fine particulate matter 2.5 (PM_2.5_) has emerged as a critical environmental risk factor for cerebrovascular diseases, contributing to an estimated 7.9 million premature deaths annually. PM_2.5_ induces cellular toxicity and is hypothesized to disrupt the blood-brain barrier (BBB), a pathological hallmark in cerebrovascular diseases such as ischemic stroke. Despite epidemiological evidence linking PM_2.5_ to increased stroke incidence, its underlying cellular mechanism driving this association is poorly understood. It remains unclear how environmentally relevant pollution concentrations affects brain endothelial function or influence stroke-related biomarkers such as the lectin-like oxLDL receptor 1 (LOX-1).

**Method:** Primary adult male human brain microvascular endothelial cells (HBMEC) were exposed to PM_2.5_ (5, 15, 75, or 300 μg/m^3^) collected from an urban environment in southern Sweden, or control. Thereafter, exposed to normoxia (21% O_2_) or hypoxia (1% O_2_) and glucose deprivation, followed by reperfusion as a model for ischemic stroke. Cell viability, oxidative stress, inflammation, BBB integrity (claudin-5, ZO-1) and LOX-1 protein expression were assessed.

**Results:** PM_2.5_ exposure induced cellular dysfunction, oxidative stress and inflammation starting at 75 μg/m^3^ PM_2.5_. Notably, decreased claudin-5 and ZO-1 protein levels and increased LOX-1 expression at concentrations as low as 15 μg/m^3^ PM_2.5_, levels commonly encountered in urban environments globally. The cellular effects of PM_2.5_ closely resembled those induced by ischemic-like injury.

**Conclusion:** These findings demonstrate dose-dependent detrimental effects of PM_2.5_ on HBMEC. The results suggest that ambient urban PM_2.5_ may act as a predisposing factor for cerebrovascular disease onset, by causing endothelial and barrier dysregulation.

**Highlights:**

- Urban PM_2.5_ dose-dependently disrupts BBB integrity in human brain endothelial cells
- PM_2.5_ induces endothelial dysfunction resembling ischemic-like injury
- Urban PM_2.5_ exposure upregulates cardiovascular disease biomarker LOX-1
- A majority of the global population are exposed to BBB-disrupting PM_2.5_ levels
- Vascular endothelial- and BBB dysfunction enhances risk for cerebrovascular disease

**Graphical abstract:** 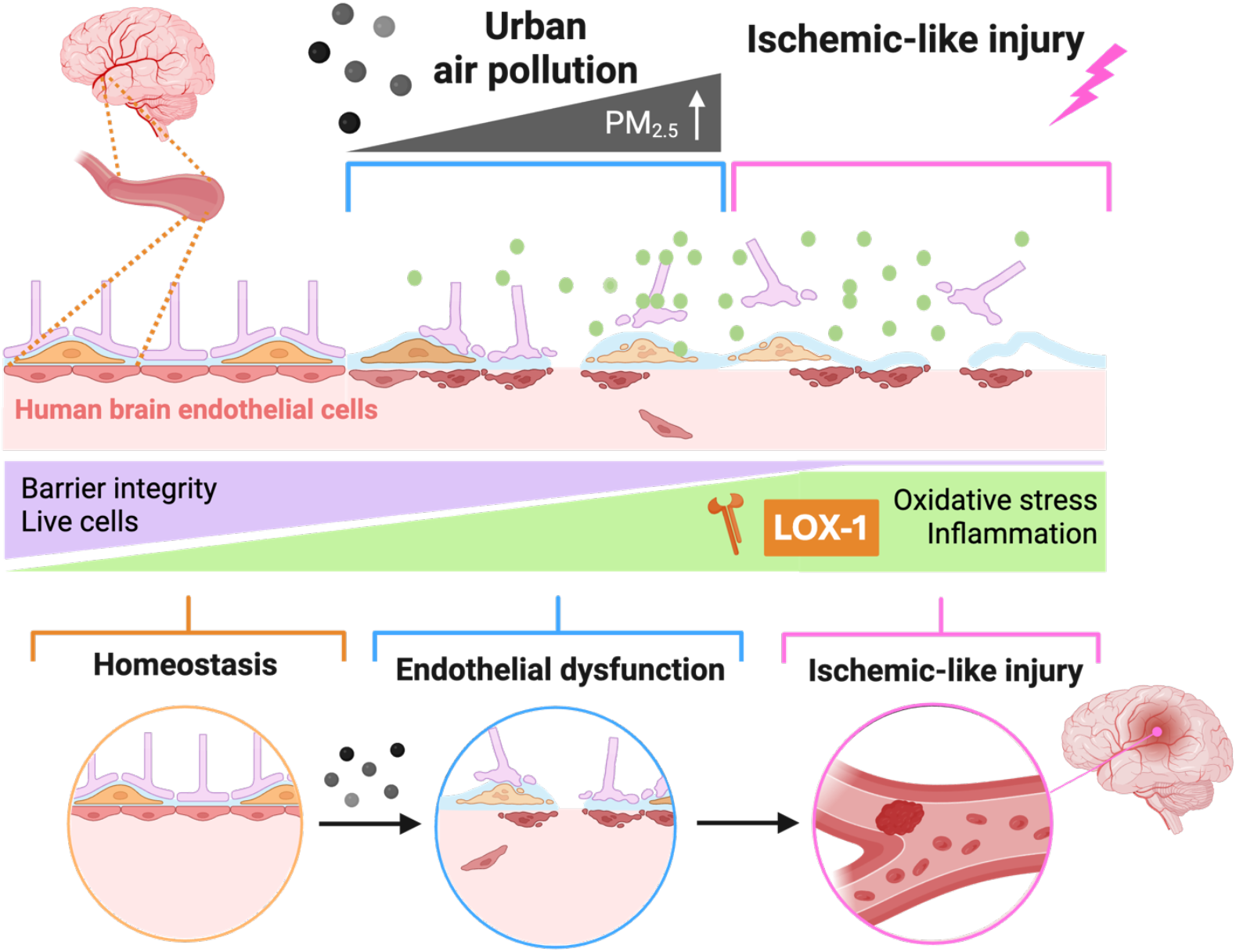

## 1. Introduction

Air pollution is widely recognized as the most significant environmental health risk globally, contributing to more than 7.9 million premature deaths annually (WHO, 2024, HEI, 2025). New safety guidelines for airborne particles with an aerodynamic diameter < 2.5 μm (PM_2.5_) were established by the World Health Organisation (WHO) in 2021 to mitigate their adverse health impacts (WHO, 2021). However, despite these updated recommendations the WHO annual maximum limit of 5 μg/m^3^ for ambient PM_2.5_ is exceeded for more than 99% of the global population (WHO, 2021, WHO, 2022). PM_2.5_ has emerged as a major risk factor for adverse health outcomes and is linked to development of respiratory, cardiovascular, cancer and neurodegenerative diseases. Strong epidemiological evidence demonstrates that elevated PM_2.5_ exposure contributes to cardiovascular diseases with high mortality and societal burden including stroke, myocardial infarction and atherosclerosis (Krittanawong et al., 2023). In addition, PM_2.5_ exposure is associated with both the onset and progression of neurodegenerative diseases such as Alzheimer’s disease that has a substantial impact on quality of life (Lee et al., 2023, Best Rogowski et al., 2025, Huang et al., 2025). Despite the widespread exposure and strong epidemiological associations with health effects relating to the brain, significant gaps remain in understanding the underlying cellular mechanisms through which PM_2.5_ contribute to disease. A major limitation in current toxicological research is the frequent use of PM_2.5_ concentrations starting around 100 μg/m^3^ (Kang et al., 2025, Chen et al., 2017, Zhu et al., 2018, Qin et al., 2023), which is far above the ambient concentrations encountered by majority of the global population. Consequently, little is known about effects of the lower, yet relevant, doses at which PM_2.5_ begins to exert harmful effects. Furthermore, it is rarely addressed how much PM_2.5_ reaches the brain upon inhalation exposure or how these particles affect brain vasculature as a large portion of pollution research focuses predominantly on the respiratory system. An additional methodological concern is that particles used in experimental studies are seldom collected from real urban environments and thereby do not accurately reflect the complex composition of air pollution particles which one is exposed to in daily life. These limitations create a substantial gap in generalizability and translational relevance of PM_2.5_ toxicity research.

Given their small aerodynamic diameter, PM_2.5_ particles can penetrate the lung epithelial barrier upon inhalation, enter the systemic bloodstream circulation and cross the blood-brain barrier (BBB) (Kang et al., 2021, Li et al., 2022). They may also reach the cerebral vasculature directly via the nasal cavity, reaching the olfactory bulb (Oberdorster et al., 2004). The BBB is a highly selective barrier which lines the cerebral vasculature and consists of endothelial cells preserved by tight junction proteins, such as claudin-5 and zonula occludens-1 (ZO-1) (Kadry et al., 2020). The barrier regulates the exchange of molecules between the blood and the brain which is crucial to maintain neural function and to protect the brain from toxins. The endothelial cells line the inner surface of blood vessels and are thereby the first point of contact between substances in the blood circulation and the BBB. Air pollution is therefore hypothesized to directly affect BBB function, and studies indicate that particle exposure can degrade tight-junction proteins and compromise barrier integrity (Oppenheim et al., 2013). PM_2.5_ also induce oxidative stress by generating reactive oxygen species (ROS) in the respiratory and vascular system, specifically in endothelial cells (Cui et al., 2015, Chen et al., 2017). Additionally, PM_2.5_ activates the immune system by promoting the release of pro-inflammatory cytokines, ultimately contributing to neuroinflammation and neurodegeneration (Gu et al., 2022, Kang et al., 2021). Endothelial dyshomeostasis and loss of barrier integrity increases barrier permeability which allows for infiltration of inflammatory cells and toxins from the blood to the brain (Takata et al., 2021). This mechanism may then augment the PM_2.5_-induced pathology through further inflammatory responses. As vascular endothelial cells represent the initial point of contact for circulating PM_2.5,_ their responses to particle exposure are highly relevant for cerebrovascular diseases.

The pathological features of air pollution on the vasculature closely resemble those observed in ischemic stroke, and accordingly, air pollution is estimated to be the second most prominent risk factor for stroke development (Collaborators, 2024). Ischemic stroke, caused by cerebral blood vessel occlusion which deprives neurons of oxygen and nutrients, leads to ROS and pro-inflammatory activation (Campbell et al., 2019, Gulke et al., 2018), causing endothelial damage and BBB degradation (Campbell et al., 2019). The lectin-like oxLDL receptor 1 (LOX-1) is upregulated in the acute phase of ischemic stroke and other cardiovascular pathologies, promoting downstream pro-inflammatory effects (Crucet et al., 2013, Arkelius et al., 2024). LOX-1 is linked to cerebrovascular endothelial dyshomeostasis and increased BBB permeability (Sawamura et al., 1997, Liang et al., 2020). Despite similarities in pathophysiology, the effects of PM_2.5_ on stroke-linked pathogenic hallmarks are yet to be studied in the brain vasculature.

Given that PM_2.5_ contributes to millions of premature deaths annually, elucidating the underlying pathological mechanism by which physiologically relevant concentrations of PM_2.5_ causes damage to the brain endothelium is crucial for disease prevention and mortality reduction. The aim of this study was to investigate the effects of urban PM_2.5_ on human brain microvascular endothelial cells (HBMEC) at concentrations reflecting WHO guidelines and real-world atmospheric exposure levels. The study examined cell viability, oxidative stress, inflammation, BBB integrity and LOX-1 expression as a stroke-linked pathogenic marker. To contextualize PM_2.5_-induced effects, the outcomes were compared to an established *in vitro* model of ischemic stroke. The effects of prior PM_2.5_ exposure on ischemic like injury outcome was also evaluated. This research addresses a critical gap in understanding how physiologically relevant concentrations of PM_2.5_ induce cerebrovascular damage at the molecular and cellular level.

## 2. Materials and methods

### 2.1 Global urban PM_2.5_ concentrations across cities as reference data for experimental concentration selection

To set environmentally realistic doses for *in vitro* exposure, data of hourly levels- and annual average PM_2.5_ concentrations, was extracted from IQAir 2024 world air quality report and IQAir live hourly PM_2.5_ measurement on website iqair.com. Annual averages were collected for Los Angeles (USA), São Paulo (Brazil), London (UK), Malmö (Sweden), Kinshasa (Democratic Republic of the Congo), Delhi (India), Beijing (China) and Sydney (Australia). Hourly fluctuation data was collected for Los Angeles, London, Malmö and Delhi at 09:00, 12:00, 17:00 and 00:00 on November 9^th^ and November 10^th^ of 2025. Different cities were selected to represent the biggest continents of the world, and to include Malmö where the studied particles were collected.

### 2.2 PM_2.5_ collection and characterisation

Urban air pollution particles were collected near a street crossing in central Malmö according to previous study protocols (Bergman et al., 2024). Briefly, PM_2.5_ was obtained using a high-volume impactor (900 l/min) (BGI900) which collected all particles < 2.5 μm on a polytetrafluorethylene membrane filter. The collected particles were extracted from the membrane using a protocol from the impactor manufacturer (BGIinc., 2008). Briefly, filters are sonicated in analysis grade methanol and the resulting dispersion is dried by vacuum evaporation. In parallel to collection, PM2.5 number size distribution and concentration were measured with high time resolution, using a scanning mobility particle sizer (SMPS) system. A section of separate filter samples was collected for chemical analysis. The PM2.5 characterization is described in detail in previous studies (Bergman et al., 2024).

The extracted particles were dispersed in ultrapure water and stored until used. Prior to treatment, the collected PM_2.5_ stock was homogenized by sonication (Branson Sonifier 250), as characterised in previous protocols (Naav et al., 2020). The stock solution was then diluted in a serial dilution with Dulbecco’s phosphate buffered saline (DPBS) to reach desired concentrations.

### 2.3 *In vitro* study design for the evaluation of dose-dependent mechanism of urban PM_2.5_ exposure

To evaluate the dose-dependent effects of PM_2.5_ on primary adult male HBMEC, cells were exposed to real-world equivalent concentrations of 5, 15, 75, or 300 μg/m^3^ ambient PM_2.5_ particles for 48h. To compare the effects of PM_2.5_ exposure with ischemic-like injury, cells were designated to either normoxia, or ischemia/reperfusion injury for 3h following the first 24h particle incubation. The plates were then reperfused by incubation under physiological conditions for 24h with corresponding dose of PM_2.5._ Cell viability and oxidative stress was assessed via cell death and ROS detection assays, together with crystal violet staining for morphological assessment. Barrier markers, inflammatory markers and LOX-1 levels was assessed through both standard western blot and in-cell western techniques. (*Figure 1*)

**Figure 1:**
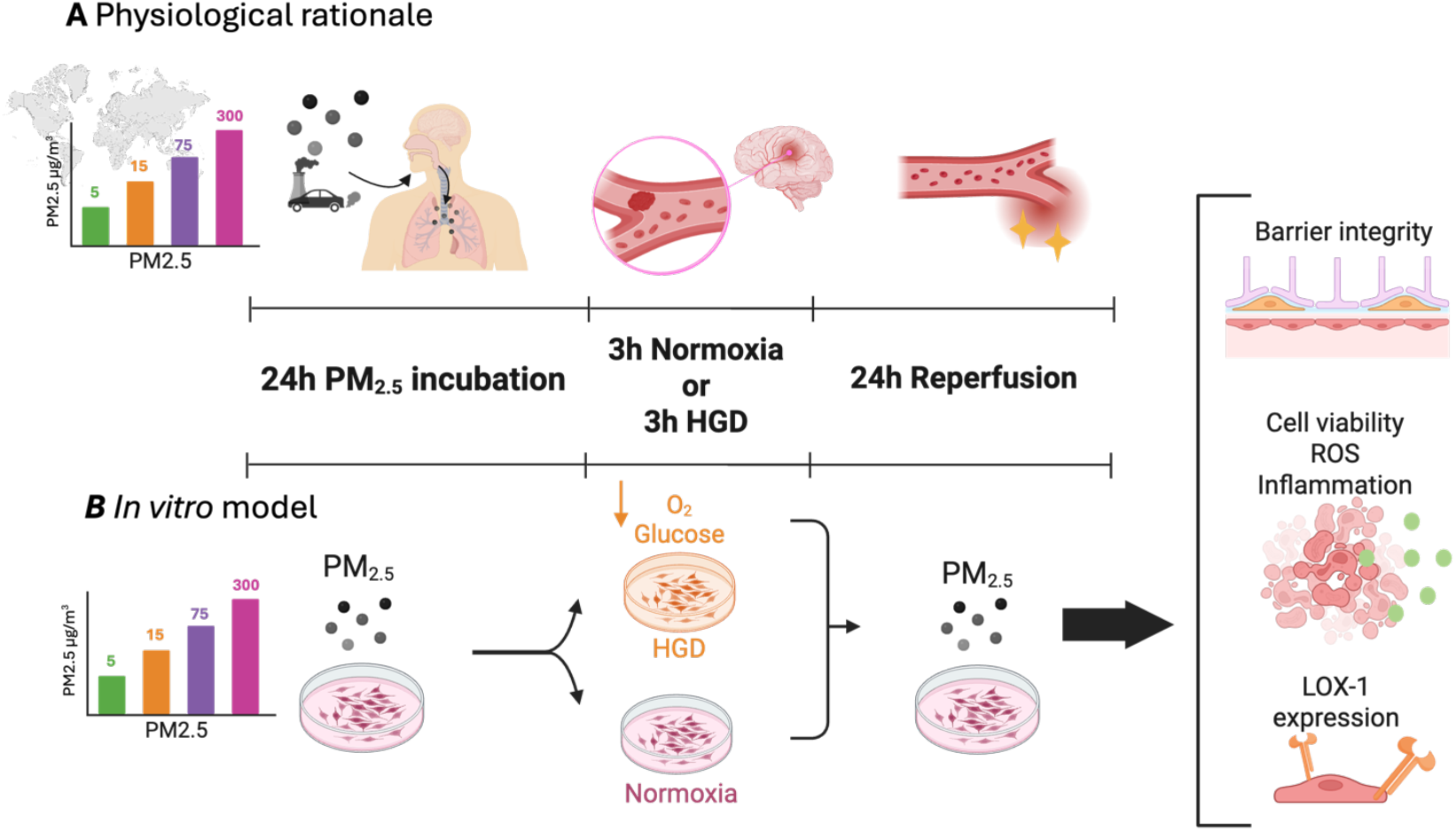
Study design for evaluating dose-dependent effects of particulate matter 2.5 (PM_2.5_) on human brain microvascular endothelial cells (HBMEC) with *in vitro* ischemic-like injury model. **A**. Physiological rationale: Ambient PM2.5 exposure is epidemiologically linked to increased ischemic stroke risk. This in vitro model simulates the real-life scenario of pre-existing PM2.5 exposure followed by ischemic stroke and subsequent reperfusion. **B**. Primary adult male HBMEC were exposed to 5, 15, 75, or 300 μg/m^3^ PM_2.5_ for 48h in total. To compare with the effects of physiological ischemic-like injury, some plates were exposed to hypoxia (1% O_2_) and glucose deprived media (HGD) for 3h after the initial 24h incubation. Following HGD or normoxia, cells were reperfused with nutrient-enriched media and incubated with PM_2.5_ at normoxic (21% O_2_) conditions as a reference for resolution of ischemia. Barrier integrity, cell viability, reactive oxygen species (ROS), inflammation and LOX-1 expression was assessed. Figure created in BioRender.

### 2.4 Cell culture and PM_2.5_ treatment

Primary adult male HBMEC were purchased from Innoprot (Spain, Catalog number: P10361, Lot number: 111224CS). Cells were cultured in endothelial cell medium (Innoprot, Catalog number: P60104) according to standard protocol. All cells were treated when 85-90% confluent at passage 6-7.

HBMEC were treated with equivalents of ambient exposure of 5, 15, 75 or 300 μg/m^3^ PM_2.5_, or 300 μg/m^3^ DPBS as vehicle control. The culturing media was replaced with nutrient enriched Dulbecco’s modified eagle medium (DMEM) (Gibco, Catalog number: 21041025) together with DPBS diluted PM2.5 solution to reach the designated concentration. Plates were incubated for 48h in a CO_2_ incubator at 37°C before collection.

The applied *in vitro* model in this study is an artificial situation which cannot be directly translated to the effects of real ambient PM_2.5_ exposure given that the calculated doses are based on estimations and assumptions of particle behaviour in physiological conditions.

To estimate PM_2.5_ doses within the vascular system, findings from investigations on how inhaled airborne nanomaterials interacts in the body were incorporated, which found that 1-2% of 60nm 99Tc aerosol particles were accumulated in the liver (Yacobi et al., 2011). Assuming the presence of other clearance pathways, an estimate of 3% crossing into the vasculature was assumed for this study. Additionally, an estimation of average volume of blood in human was defined to 5L. These factors were utilized to calculate the appropriate PM_2.5_ concentrations representative of atmospheric exposure to PM_2.5_ levels of 5, 15, 75, and 300 μg/m^3^. As a result, the final concentrations of PM_2.5_ within the cultures were 0.33, 0.99, 4.95, and 19.8 μg/L corresponding to the atmospheric concentrations of 5, 15, 75, and 300 μg/m^3^ respectively. However, translocation rates reported in the literature show considerable variability, and our applied concentrations may represent an upper-bound estimate of physiological endothelial exposure.

Furthermore, characterization of the applied particles reveals a large ultrafine fraction (aerodynamic diameter of 0.1μm) which may exhibit different translocation kinetics than typical PM_2.5_. Given this size heterogeneity and the current uncertainties in translocation dynamics for real-world particle mixtures, we acknowledge that precise translation of ambient exposure to *in vitro* endothelial concentrations is difficult to estimate the exact physiological distribution.

### 2.5 *In vitro* ischemia/reperfusion injury

*In vitro* ischemia/reperfusion injury (HGD) was modelled based on methods of previous studies (Arkelius et al., 2024, Wendt and Gonzales, 2023). After initial 24h PM_2.5_ exposure, cells designated for HGD were washed with DPBS and had their media replaced with DMEM deprived of glucose, glutamine, phenol red and sodium (Gibco, Catalog number: A1443001). The plates were then incubated in a hypoxic chamber (Stemcell technologies) flushed with a gas mixture containing 1% O_2_, 5% CO_2_ and nitrogen balance, in a CO_2_ incubator for 3h. To simulate oxygen and nutrient reperfusion following ischemic stroke resolution, the plates were removed from the hypoxia chamber and the glucose deprived DMEM was replaced with nutrient enriched DMEM. Designated PM_2.5_ concentrations were added to the plates which were then incubated in a CO_2_ incubator for another 24h before analysis.

For measurement of in-cell Western, cell death, oxidative stress, and crystal violet staining, cells were seeded in 96-well plates or on glass-slides to reach a density of 1-2 x 10^4^ cells and treated as described in section 2.4 and 2.5.

### 2.6 In-cell Western

Protein expression of tight junction proteins claudin-5 and ZO-1 was assessed with an in-cell Western kit according to manufacturer’s instructions (LicorBio, Catalog number: NC2225468). 926-42094). HBMEC treated in 96-well plates were fixed with 4% formaldehyde and permeabilized with 0.1% Triton X-100. Cells were blocked with Intercept Blocking Buffer (LicorBio, Catalog number: 927-60001) and incubated with primary antibody; claudin-5 (1:200, ThermoScientific, Catalog number: 34-1600) and ZO-1 (1:200, ThermoScientific, Catalog number: 61-7300) overnight at 4°C. The cells were then washed and incubated with secondary antibody; anti-rabbit (1:1000, Li-CorBio, Catalog number: 926-3221) and CellTag520 stain (LicorBio, Catalog number: 926-41094) for 1h at RT to detect claudin-5 and cell nuclei. Fluorescent signal was measured using BMG Labtech ClarioStar Plus. Total claudin-5 and ZO-1 protein signal was normalized to the corresponding CellTag520 stain signal. (Wendt and Gonzales, 2023)

### 2.7 Cell death and ROS assay

Treated HBMEC were washed with DPBS to remove excess particles. To evaluate cell death, CyQUANT cell proliferation assay was used (Invitrogen, Catalog number: C35011). A cell-permeant nucleic acid stain and a non-permeant background suppressor was incubated with the treated cells for 1h at 37°C. The plates were then imaged with an excitation and emission wavelength of 480- and 535nm respectively and a gain of 3000 (BMG Labtech, ClarioStar Plus). To assess ROS levels, dichorofluorescein diacetate (DCHF-DA) (Sigma-Aldrich, Catalog number: 35845) was dissolved in dimethyl sulfoxide (DMSO) and diluted in DPBS to a 20 μM working solution, which was then incubated with the cells for 30 min at 37°C in the dark. Following incubation, cells were washed and imaged (BMG Labtech, ClarioStar Plus) with an excitation wavelength of 485nm, emission wavelength of 520nm and gain of 2000.

### 2.8 Crystal violet staining

HBMEC morphology was visualized with crystal violet staining. Cells were seeded and treated on coverslips as previously described (2.4), washed, and fixed with 4% formaldehyde and incubated with crystal violet solution for 20 min at RT. The coverslips with stained cells were mounted on glass slides and imaged (Olympius Slideview VS200). Staining protocol was developed from previous studies (Wendt et al., 2021).

Cellular lengths were quantified from crystal violet images of using FIJI (ImageJ2 v2.16.0/1.54p). Prior to measurement, all images were calibrated to the spatial resolution of Olympus Slideview using the pixel size embedded in the acquisition metadata. To ensure accurate cell boundary visualization, raw images were converted to 8-bit grayscale and minimally adjusted for contrast without altering relative signal intensities. Individual cells were selected for analysis only if they were isolated, fully contained within the field of view, and not undergoing division or detachment. For each cell, its longitudinal axis was traced manually using the segmented line tool, placing sequential vertices along the central axis of the cell from one end of the membrane to the other. This approach ensured that curved or irregular cell morphologies were accurately represented. Upon completing each trace, the total path length was quantified which returned cell length in micrometers based on the previously calibrated scale.

### 2.9 Protein extraction and concentration determination

Treated cells were collected and combined with denaturing cell extraction buffer (ThermoScientific, Catalog number: FNN0091) containing protease- and phosphatase inhibitor (Sigma-Alrich, Catalog number: P8340, P2850). The samples were homogenized and centrifuged to collect cell lysate. Total protein concentration was determined using a Bio-Rad protein assay (Bio-Rad, Catalog number: 5000006), measuring absorbance at 595nm on a microplate reader. Final protein concentrations were calculated from the mean absorbance value and corrected against a BSA standard curve.

### 2.10 Western Blot

Western Blot was run to visualize changes in protein expression of pro-inflammatory cytokine interleukin-6 (IL-6) and LOX-1 (Orset et al., 2021). Denaturing cell extraction buffer, Laemmli reagent (Bio-Rad, Catalog number: 1610737EDU) and 2-mercaptoethanol was boiled with the protein samples. Proteins were separated by SDS-PAGE on a 4-20% Tris-Glycine gel and electrotransfered to a nitrocellulouse membrane (Bio-Rad, Catalog number: 1620112). The membrane was then blocked; 5% BSA in T-TBS for IL-6, or Starting block for LOX-1 (ThermoScientific, Catalog number: 37538). Following, the membrane was incubated with primary antibody; IL-6 (1:250, Abcam, Catalog number: Ab6672) or LOX-1 (1:1000, Abcam, Catalog number: Ab60178) overnight at 4°C. Thereafter, the membrane was washed and incubated with secondary antibody; anti-rabbit (1:2000, Cell Signaling Technology, Catalog number: 7074) for 1h at RT and imaged using SuperSignal Dura kit (ThermoScientific, Catalog number: 34075) in FujiFilm LAS-1000 image analyzer. Subsequently, the membranes were stripped (ThermoScientidic, Catalog number: 21059) and probed for β-actin (1:50,000, Sigma-Aldrich, Catalog number: A3854). Signal quantification was performed using FIJI and normalized to β-actin signal as loading control. The individually loading-normalized values were then represented relative to the corresponding vehicle-treated control.

### 2.11 Statistical analysis

All primary HBMEC samples are replicates originating from one donor, where “n” corresponds to number of technical replicates independently repeated at least twice. Statistical analysis was performed using GraphPad Prism and all data is presented as mean ± SD. Multiple comparisons were assessed with one-way or two-way non-parametric ANOVA test (Kruskal-Wallis) and direct comparisons between two groups was assessed with Mann-Whitney test. P < 0.05 was considered statistically significant.

## 3. Results

### 3.1 Air PM_2.5_ concentrations exceeds WHO recommendation in multiple cities globally

Of the assessed cities, Sydney had the lowest average PM_2.5_ concentration of 4.3 μg/m^3^ in 2024, followed by Malmö (7 μg/m^3^), London (7.8 μg/m^3^), Los Angeles (10.1 μg/m^3^), São Paulo (15.9 μg/m^3^), Beijing (30.9 μg/m^3^), Kinshasa (58.2 μg/m^3^) and lastly Delhi with the highest average of 108.3 μg/m^3^. The WHO recommended maximum annual PM_2.5_ exposure of 5 μg/m^3^ is exceeded in all compared cities except for Sydney. Comparison of air pollution concentrations on November 9^th^ or 10^th^ of 2025 in Los Angeles, London, Malmö and Delhi shows that PM_2.5_ concentrations fluctuate throughout the day in all cities. PM_2.5_ in Los Angeles ranged between 12.6 and 19.6 μg/m^3^, 6.0-15.3 μg/m^3^ in London, 8.8-17.9 μg/m^3^ in Malmö and 203-462 in Delhi. (*Figure 2*)

**Figure 2:**
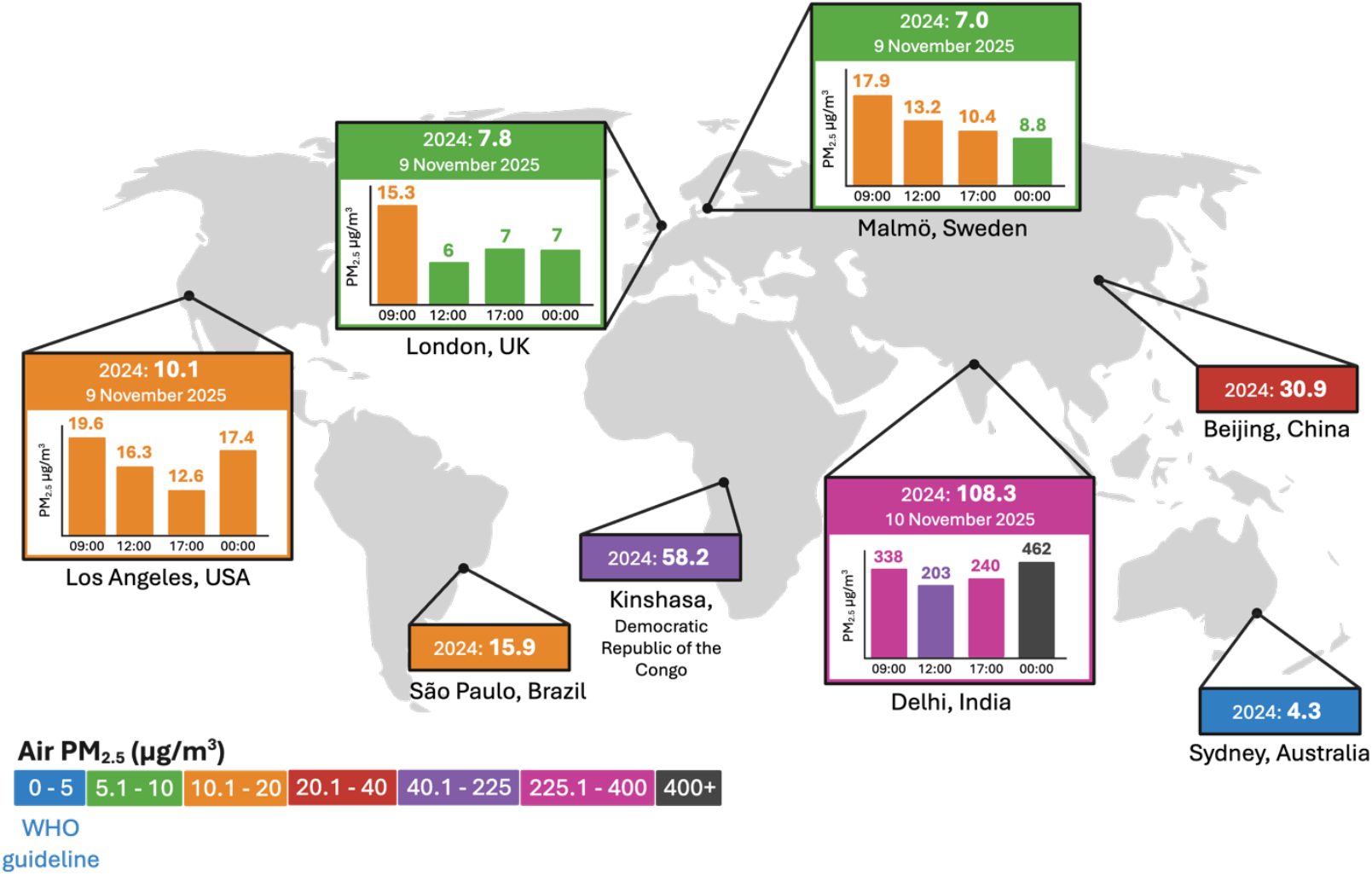
Comparison of atmospheric particulate matter 2.5 (PM_2.5_) concentrations worldwide. Measurements of average- and daily air PM_2.5_ levels in Los Angeles, São Paulo, Kinshasa, London, Malmö, Delhi, Beijing and Sydney according to IQAir 2024 world air quality report and IQAir live PM_2.5_ measurement. Comparison shows that average PM_2.5_ concentrations of 2024 differs drastically between cities in different areas of the world, including Sydney (4.3 μg/m^3^) which is the only assessed city within the recommended WHO guideline, Malmö (7.0 μg/m^3^), London (7.8 μg/m^3^), Los Angeles (10.1 μg/m^3^), São Paulo (15.9 μg/m^3^), Beijing (30.9 μg/m^3^), Kinshasa (58.2 μg/m^3^) and Delhi with the highest PM_2.5_ average of 108.3 μg/m^3^. Data of differences in PM_2.5_ concentrations during the day fluctuated in all compared cities and ranged between 12.6-19.6 μg/m^3^ in Los Angeles, 6.0-15.3 μg/m^3^ in London, 8.8-17.9 μg/m^3^ in Malmö and 203-406 μg/m^3^ in Delhi on November 9^th^ or 10^th^ of 2025. Figure created in BioRender with data from iqair.com. Map lines delineate study areas and do not necessarily depict accepted national boundaries.

### 3.2 PM_2.5_ reduces expression of BBB tight-junction proteins claudin-5 and ZO-1 in HBMEC

Total claudin-5 protein levels decreased dose-dependently with higher PM_2.5_ concentration in HBMEC with in-cell western assessment from concentrations ≥15μg/m^3^ compared to vehicle (*Figure 3A*). There was no difference in protein expression between PM_2.5_ doses when exposed to ischemic-like injury; however, HGD exposed cells had lower protein expression in comparison to cells treated with only PM_2.5_. Similarly, ZO-1 expression declined from exposure to 15- and 300μg/m^3^ PM_2.5_ (*Figure 3B)*. Kruskal-Wallis multiple comparison analysis of ZO-1 expression also calculated p = 0.06 between vehicle and 5μg/m^3^ and p = 0.1017 between vehicle and 75μg/m^3^.

**Figure 3:**
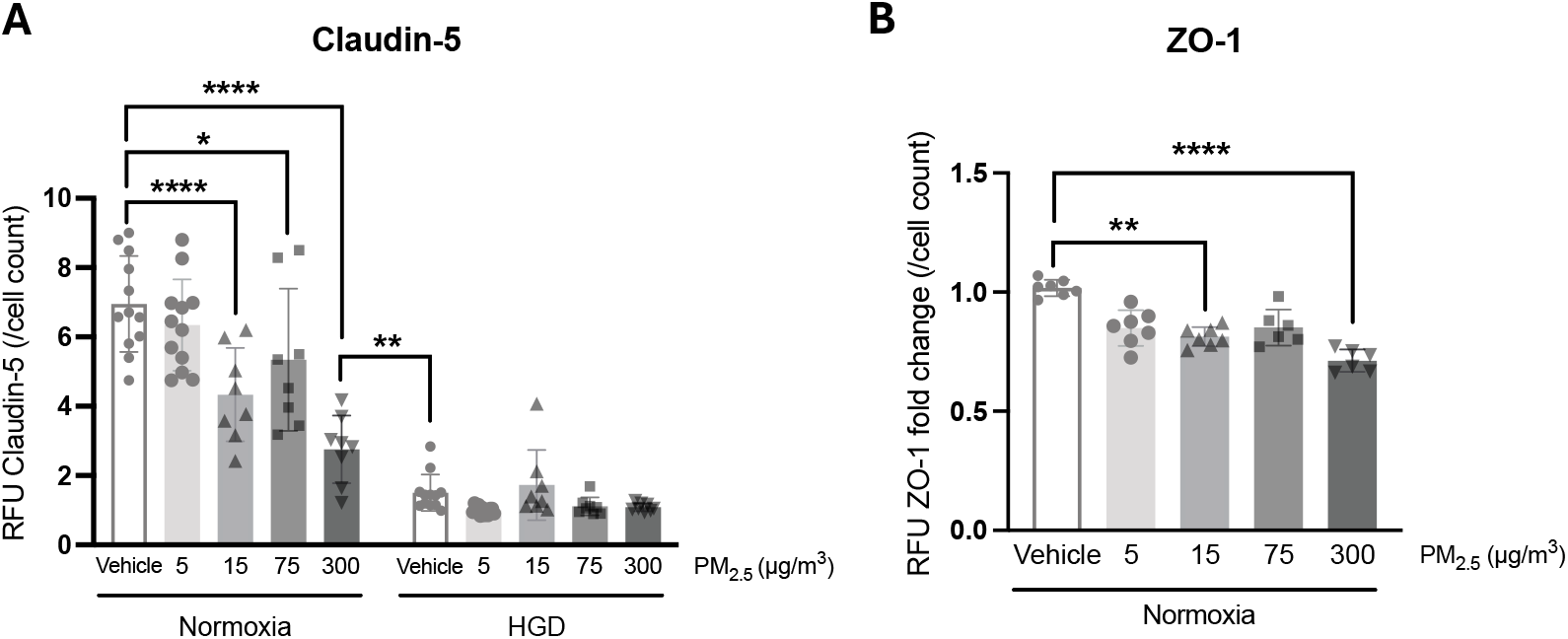
Decreased expression of tight-junction proteins claudin-5 and zonula occludens-1 (ZO-1) in human brain microvascular endothelial cells (HBMEC) with exposure to particulate matter 2.5 (PM_2.5_). HBMEC exposed to PM_2.5_ (5, 15, 75, or 300 μg/m^3^) for 24h and 3h normoxia- or hypoxia with glucose deprivation (HGD) followed by 24h reperfusion. **A**. In-cell Western analysis of total claudin-5 protein expression reveals a dose-dependent decreased protein expression in normoxic cells from exposure to 15 μg/m^3^ and higher. Claudin-5 levels remained constant in HGD treated cells across all PM_2.5_ concentrations, however significantly lower in comparison to normoxic HBMEC. (n=12 technical replicates for vehicle and 5, n=8 technical replicates for 15, 75 and 300). **B**. In-cell Western analysis of total ZO-1 expression fold-change shows decreased protein expression from exposure to 15 μg/m^3^ and 300 μg/m^3^. (n=6-7 technical replicates). Data presented as mean +-SD. Statistical significance assessed through Kruskal-Wallis test within treatment groups (Normoxia/HGD) and Mann-Whitney test between groups with different treatment (300 normoxia / vehicle HGD). *p<0.05, **p<0.01, ****p<0.0001

### 3.3 PM_2.5_ disrupts HBMEC viability and increases oxidative stress dose-dependently

Live cell count of PM_2.5_ exposed HBMEC decreased dose-dependently with ascending PM_2.5_ dose, from concentrations ≥75μg/m^3^ compared to vehicle-treated controls (*Figure 4A*). HGD exposure significantly reduced cell viability compared to normoxic groups but showed no additional effects between exposure to different PM_2.5_ concentrations. Similarly, data shows a significant dose-dependent increase in ROS levels from concentrations ≥75μg/m^3^ compared to vehicle (*Figure 4B*). HGD treatment elevated ROS levels in comparison to normoxic groups but remained stable across all PM_2.5_ concentrations. Crystal violet staining and analysis of cellular length visualizes that HBMEC treated with ≥15μg/m^3^ PM_2.5_ exhibited elongated morphology in comparison to vehicle (*Figures 4C-D*). Additionally, HBMECs appeared to have erratic cellular borders which gradually extended towards neighbouring cells when exposed to higher air pollution concentrations.

**Figure 4:**
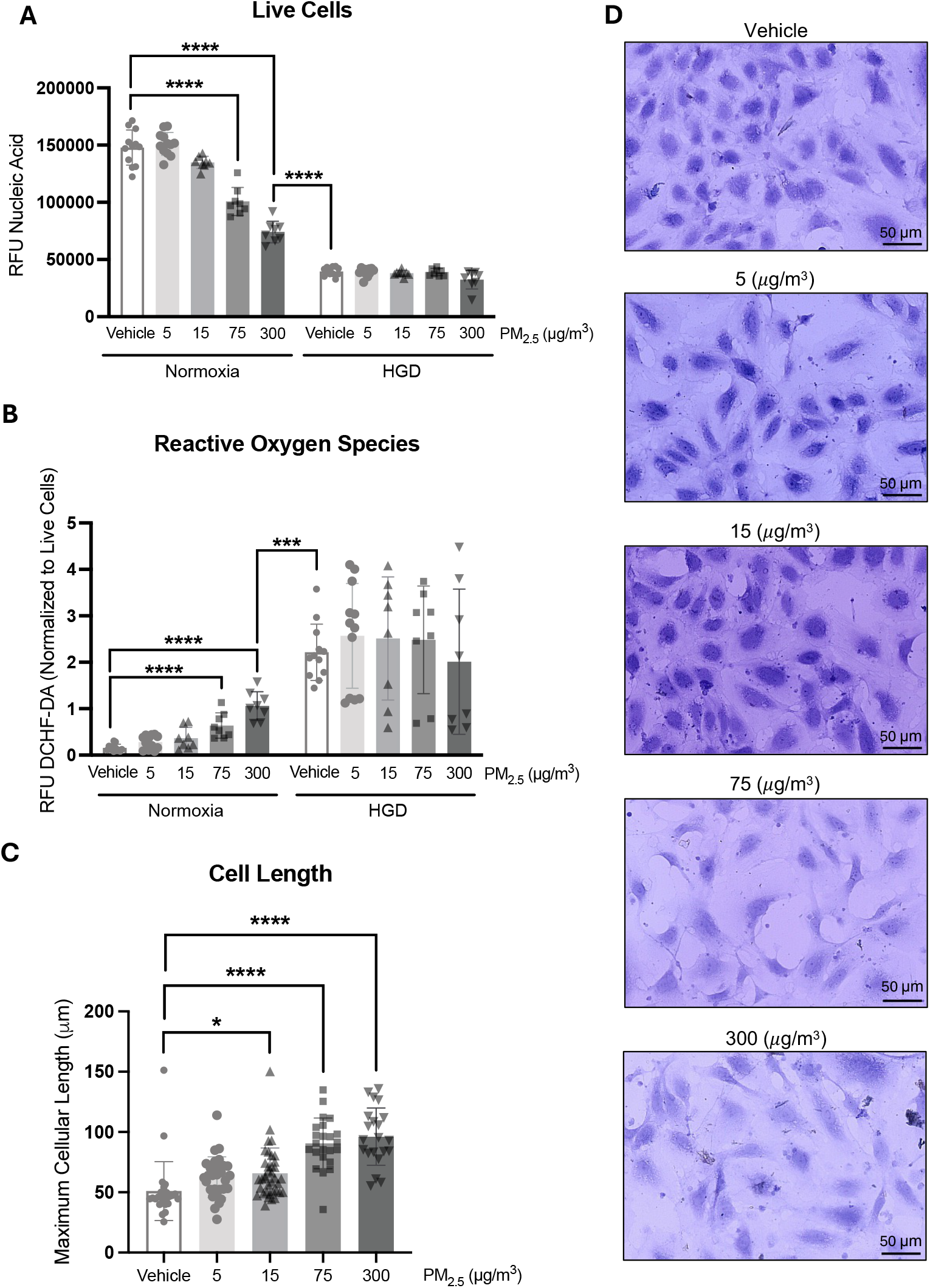
Differentiated live cell count, oxidative stress and cellular morphology of human brain microvascular endothelial cells (HBMEC) exposed to particulate matter 2.5 (PM_2.5_) during normoxia and ischemia. Adult male HBMEC were exposed to vehicle or PM_2.5_ (5, 15, 75, or 300 μg/m^3^) for 24h and incubated for 3h in normoxia- or hypoxia and glucose deprivation (HGD) followed by 24h reperfusion. **A**. Live cell count (CyQUANT nuclear stain) decreased when exposed to ≥75 μg/m^3^ PM_2.5_ compared to vehicle. HGD treatment reduced live cell count compared to normoxia but did not differ between particle treated groups. **B**. Reactive oxygen species (ROS) signal (DCHF-DA) normalized to live cell count. Relative ROS levels increased dose-dependently with PM_2.5_ concentration, with significant increase observed at PM_2.5_ ≥75 μg/m^3^, in comparison to normoxia vehicle. ROS levels were uniformly elevated following HGD across all doses in comparison to normoxia vehicle and significantly higher than untreated HBMEC. (n=12 technical replicates for vehicle and 5, n=8 technical replicates for 15, 75 and 300) **C**. Analysis of crystal violet-stained HBMEC shows a longer maximum cellular length when treated with ≥15 μg/m^3^ PM_2.5_. (n=21-37 individual cells) **D**. Representative images of crystal violet-stained HBMEC visualizing a differentiated morphology in cells treated with higher PM_2.5_ concentration, where cells appear more elongated and expanding towards neighbouring cells. Data presented as mean ± SD. Statistical significance assessed through Kruskal-Wallis test within treatment groups (Normoxia/HGD) and Mann-Whitney test between groups with different treatment (300 normoxia / vehicle HGD). *p<0.05. ***p<0.001. ****p<0.0001.

### 3.4 Increased IL-6 and LOX-1 protein expression from PM_2.5_ exposure and *in vitro* ischemia/reperfusion injury in HBMEC

Western Blot signal quantification of pro-inflammatory cytokine IL-6 visualized no difference in 25kDa isoform between any groups (*Figures 5A-B*). However, there was a dose-dependent increase in the 17kDa isoform with PM_2.5_ exposure ≥75μg/m^3^ and HGD treatment compared to vehicle-treated control cells (*Figures 5A & C*). LOX-1 protein expression increased from exposure to PM_2.5_ ≥15μg/m^3^ and from ischemic like injury (*Figures 5D-E*).

**Figure 5:**
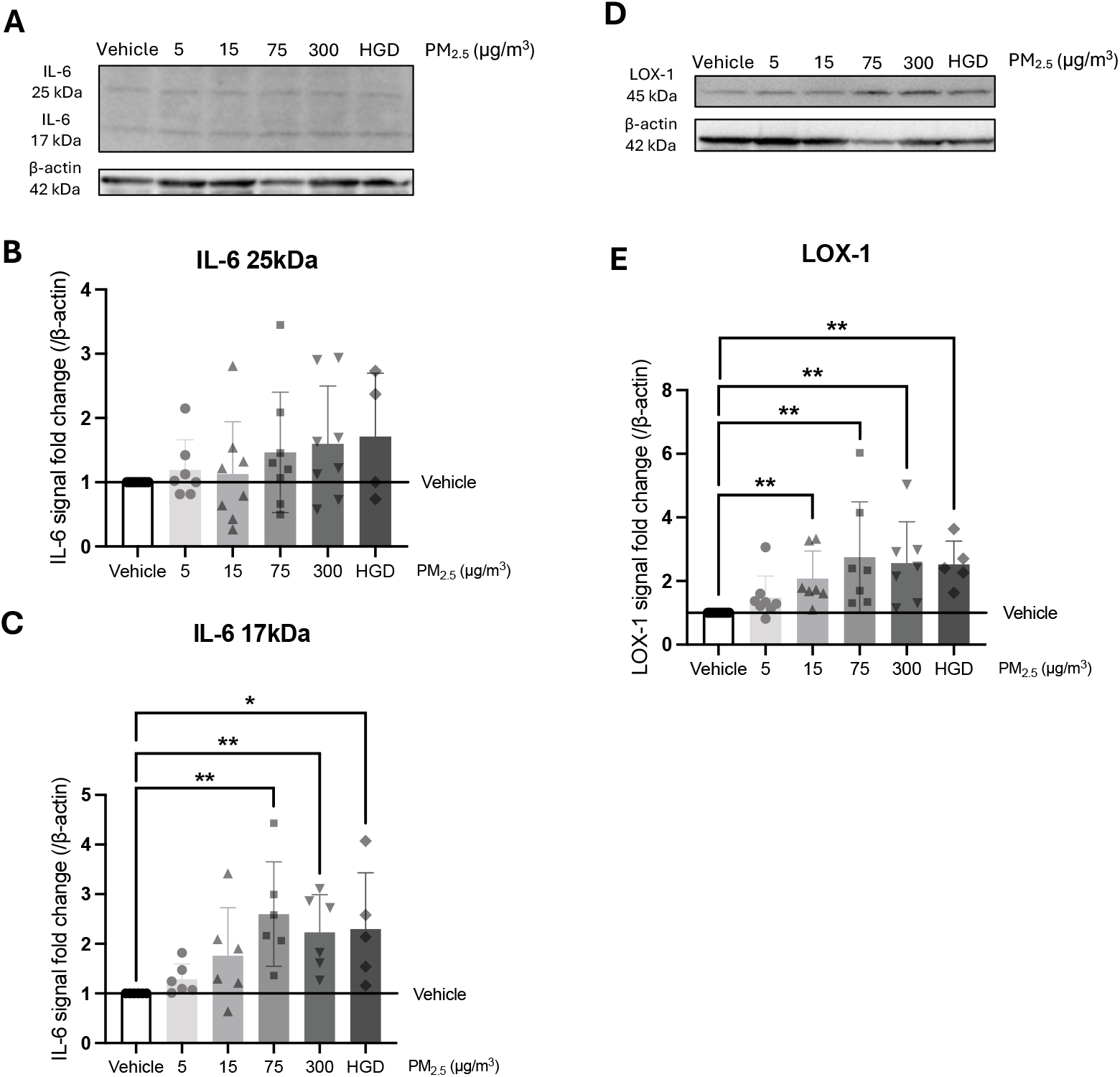
Dose-dependent increase in IL-6 and LOX-1 protein levels in human brain microvascular endothelial cells (HBMEC) exposed to particulate matter 2.5 (PM_2.5_). Western Blot assessment of adult male HBMEC exposed to vehicle, 5, 15, 75, or 300 μg/m^3^ PM_2.5_ during normoxia or ischemic-like injury with hypoxia, glucose deprivation and reperfusion (HGD). **A**. Representative Western Blot image of IL-6 and β-actin band migration. **B**. Signal quantification of 25kDa IL-6 shows no difference between PM_2.5_ exposure or HGD treated group. **C**. Signal quantification of 17kDa IL-6 shows dose-dependency with higher IL-6 expression from higher PM_2.5_ exposure, with significant increase ≥75 μg/m^3^ and from HGD treatment compared to vehicle. **D**. Representative Western Blot image of LOX-1 and β-actin. **E**. Signal quantification of LOX-1 displays a dose-dependent increase in LOX-1 with exposure to ≥15 μg/m^3^ PM_2.5_ or HGD. (n=4-7 technical replicates). Data presented as mean +-SD. Statistical significance assessed by Kruskal-Wallis test. *p<0.05, **p<0.01.

## 4. Discussion

The present study investigated the concentration-dependent effects of real-life urban PM_2.5_ exposure on HBMEC, focusing on key biological mechanisms implicated in the pathogenesis of cerebrovascular diseases. Endothelial cell viability, claudin-5 and ZO-1 protein expression of the BBB, oxidative stress and inflammation were evaluated in both normoxic- and ischemic conditions. Changes in LOX-1 expression following PM_2.5_ exposure was also assessed. Characterised PM_2.5_ particles collected from real urban environments were applied at concentrations reflective of amounts which may be absorbed and reach the cerebrovasculature via the blood circulation from environmentally relevant concentrations to higher exposure levels, thereby reflecting multiple real-world human exposure scenarios. The findings demonstrate that PM_2.5_ induces clear dose-dependent alterations on the studied parameters, with notable effects on cellular morphology, claudin-5, ZO-1 and LOX-1 expression already at 15 μg/m^3^, representative of conditions experienced by majority of the global population. Thus, this study addresses a critical gap in PM_2.5_ research by examining environmentally and physiologically relevant concentrations at- and above the WHO’s safety limit of 5 μg/m^3^. This comprehensive assessment on human tissue thereby reveals the mechanistic pathways through which low-level PM_2.5_ exposure may contribute to systemic and cerebrovascular disease in humans.

The present study concerns key components of the BBB, including tight-junctions and endothelial cells using human *in vitro* model. The loss of tight-junction proteins claudin-5, ZO-1 and reduced cell viability following PM_2.5_ exposure was similar to the decline observed after ischemic-like injury, however, ischemia resulted in a larger loss of protein and live cells compared to only PM_2.5_. Importantly, when ischemia was induced, no difference emerged between PM_2.5_ concentrations, suggesting that air pollution triggers similar but less severe pathological mechanism and does not exacerbate ischemia damage once injury is initiated. Because endothelial integrity is crucial for maintaining vascular homeostasis and BBB function, its disruption increases infiltration vulnerability to cerebrovascular and neurodegenerative pathology. In stroke, BBB breakdown enables rapid influx of inflammatory cells and toxins driving neuroinflammation and brain damage (Yang et al., 2019). In Alzheimer’s disease, a slower but progressive increase in barrier permeability contributes to pathological progression (van de Haar et al., 2016a, van de Haar et al., 2016b, Fiala et al., 2002). Since air pollution is associated with both stroke and neurodegenerative diseases (Kim et al., 2025, Dorsey et al., 2025), the early tight-junction breakdown and loss of endothelial barrier may be a shared mechanism by which PM_2.5_ contributes to multiple cerebrovascular diseases. Additionally, the mechanism causes influx of pollution particles, directly causing neural damage (Kang et al., 2021). These findings identify PM_2.5_ exposure as a risk factor for stroke induction, causing BBB dysfunction through tight-junction and endothelial damage which makes the BBB more susceptible to develop cerebrovascular diseases.

PM_2.5_ exposure further reduced HBMEC viability and barrier integrity by concentration dependent increase in intracellular ROS and IL-6 release, indicating activation of oxidative stress and inflammation. These findings of ROS and cytokine activation align with previous studies, reporting increased oxidative stress and reduced cell viability in human microvascular endothelial cells exposed to 200 μg/mL PM_2.5_ and elevated IL-6 expression in lung epithelial-endothelial co-culture models exposed to 60 μg/mL PM_2.5_ (Chen et al., 2017, Wang et al., 2019), which would correspond to around 2.0 × 10^8^ and 6.0 × 10^7^ μg/m^3^ of airborne exposure, respectively. Thus, this study observes similar effects from lower PM_2.5_ concentrations, in mechanisms which are central to the development of pollution-associated cardiovascular and cerebrovascular diseases. This study also concludes that LOX-1 expression increases from PM_2.5_ exposure and is equally expressed from ischemic-like injury. This has been seen in the brain microvasculature of high-fat diet and apolipoprotein knock out rodents when exposed to 100 μg/m^3^ diesel and gasoline exhaust (Suwannasual et al., 2018, Lucero et al., 2017), but has not been studied with PM_2.5_ in healthy rodents, to our knowledge. Thereby, this study is unique in observing upregulation of a stroke-linked biomarker when exposed to as low as 15 μg/m^3^ PM_2.5_. Similar rodent research links diesel and gasoline particle exposure with upregulation of matrix metalloproteinases (MMPs) (Oppenheim et al., 2013), enzymes capable of degrading extracellular matrix components. Mechanistically, LOX-1 can directly upregulate MMP-9 (Li et al., 2003), a matrix metalloproteinase worsening outcome in multiple neurovascular diseases as it contributes to BBB breakdown (Rempe et al., 2016, Inzitari et al., 2013). Specifically, research point towards that MMP-9 degrades both claudin-5 and ZO-1 (Feng et al., 2011). This aligns with the findings of this study, reporting upregulation of LOX-1 and downregulation of claudin-5 and ZO-1 with 15 μg/m^3^ PM_2.5_, suggesting that LOX-1 mediates barrier damage from air pollution via initiating MMP production. The receptor is associated with multiple cardiovascular pathologies as it binds- and internalizes circulating oxidised low-density lipoproteins in the vascular endothelium, forming atherosclerotic plaques that can evolve into thrombi and cause both stroke and myocardial infarction (Sanchez-Leon et al., 2024). Taken together, this study suggests that LOX-1 is a key component in PM_2.5_ induced endothelial damage as the receptor has downstream effects for multiple PM_2.5_ associated diseases.

Since both BBB damage and LOX-1 upregulation is seen from 48h exposure to only 15 μg/m^3^ PM_2.5_, it is of importance to raise the question of how air pollution levels strategically can be managed to prevent disease. While the key solution is to limit air pollution exposure, it is of interest to develop a safe and effective pharmaceutical candidate to mitigate PM_2.5_ induced damage when atmospheric exposure cannot be managed within the safety guidelines. Clinically, LOX-1 is mentioned as a promising therapeutic target in multiple cardiovascular diseases (Markstad et al., 2019, Arkelius et al., 2024, Schiopu et al., 2023), and inhibition of the receptor have showed promising results in phase 1 clinical trial on type 2 diabetes (Vavere et al., 2023). Given the early effects which PM_2.5_ has on LOX-1 in this study, receptor inhibition may be a key in mitigating endothelial- and BBB dysfunction to prevent disease. It is thereby of further interest to investigate the role of LOX-1 in PM_2.5_ induced pathogenesis, both relating to stroke and other linked diseases. Although, reducing air pollution levels is the most direct approach to avoid health effects.

Global comparison of PM_2.5_ concentrations show substantial variations across regions with levels fluctuating within urban areas over the course of the day. Although cities such as London and Malmö report annual averages around 7 μg/m^3^ air PM_2.5_ in 2024, residents are regularly exposed to over 15 μg/m^3^ during morning rush hours. This indicates that residents of most surveyed cities intermittently encounter pollution concentrations capable of inducing brain endothelial damage by tight junction degradation. The ability to control and define exposure concentrations in vitro is critical, as it allows identification of dose–response relationships and cellular thresholds for toxicity that cannot be resolved in population-based studies. While epidemiological studies associate PM_2.5_ exposure with increased disease risk, most are limited by cross-sectional designs and residual confounding related to lifestyle factors and regional variability of particle composition between populations, which cannot be disregarded as potential causal components to disease development. The present findings therefore provide mechanistic insight by demonstrating that PM_2.5_ directly impairs human brain endothelial function. Integrating controlled, concentration-defined experimental models with epidemiological evidence is essential for strengthening causal inference and for informing exposure limits, risk assessment, and intervention strategies aimed at reducing the cerebrovascular health impacts of air pollution. However, it is important to acknowledge that *in vitro* modelling, like epidemiological studies, cannot be directly translated to real physiological effects of air pollution. Our model represents an artificial system with concentration dosing based on estimations of physiological distribution of inhaled PM_2.5_. Additionally, PM_2.5_ may change composition when translocated across the blood-air barrier in the alveoli and in the blood circulation as organic material may not be translocated across the barrier (Soares et al., 2025). Organic components may desorb, degrade, or fail to cross biological barriers, while lipophilic chemicals and inorganic ions likely constitute a larger fraction of systemically available material. This represents a significant limitation of our study: by directly exposing human endothelial cells to collected urban PM_2.5,_ we do not replicate the selective translocation and transformation that occurs *in vivo*. While the precise composition of particles and chemicals reaching the cerebral endothelium remains largely unknown, studies suggest that water-soluble ions and metal components may mediate many health-related effects of air pollution (Wang et al., 2020, Yan et al., 2023). Although, the individual physiological effects of different PM_2.5_ components remains unknown, the oxidative stress observed in this study may be caused by inorganic species such as nitrates and other ions in the sample with oxidative properties. Nevertheless, our use of human endothelial cells provides mechanistic insights into potential cellular responses that are difficult to evaluate through alternative approaches.

Specifically studying and characterising PM_2.5_ particles from urban environments makes the study design attributable to a larger population and not only to specific exposure areas. The collection and characterisation of PM_2.5_ in this study revealed that the majority of urban particles had a diameter of less than 0.1μm, thus, ultrafine particulate matter (Bergman et al., 2024). Given that ultrafine particles can penetrate even deeper in the respiratory tract, enter the circulation, reach the cerebrovascular tract, and excert BBB disrupting effects, they are a big danger to global health. Thus, hypothesizing that ultrafine particulate matter exerts the molecular effects seen in this study. These results strengthen the current discussion in using ultrafine particulate matter as exposure metric in opposition to the current PM_2.5_ (Morawska et al., 2024), as the smaller diameter penetrates deeper into physiological structures. The European Union Air Quality Directive (2024/2881) also emphasizes ultrafines as an emerging pollutant of concern (EuropeanUnion, 2024).

In conclusion, this study highlights the critical role of environmental concentrations of PM_2.5_ in mediating cellular dysfunction through impairing cell viability, inducing oxidative stress, inflammation and barrier disruption, and uniquely confirms upregulation of the scavenger receptor LOX-1 in response to air pollution exposure. These findings emphasize the cerebrovascular vulnerability to air pollution and underscore the relevance of studying real urban PM_2.5_ at a broad spectrum of doses in multiple diseases which are connected to air pollution. Thus, providing valuable insights into the underlying mechanisms by which PM_2.5_ contributes to the development of cerebrovascular disease.

## Declaration of competing interests

The authors declare that they have no competing financial or personal interests that could have appeared to influence the work reported in this paper.

## CRediT authorship contribution

**Elisabet Andersson**: Writing – original draft, writing – review & editing, data curation, formal analysis, investigation, methodology, visualization, validation. **Trevor Wendt**: Writing – review & editing, formal analysis, investigation, methodology, visualization, validation. **Fanny Bergman**: Writing – review & editing, investigation, methodology, resources. **Christina Isaxon**: Writing – review & editing, investigation, methodology, resources. **Saema Ansar**: Writing – review & editing, conceptualization, funding acquisition, methodology, project administration, resources, supervision.

## Acknowledgements and funding sources

This work was supported by the Swedish Research Council (VR 2023-030004) (AS), Thure Carlssons Foundation (AS) and AFA (160226) (IC).

## Notes

### Competing Interest Statement

The authors have declared no competing interest.

### Summary of Updates

Manuscript is updated to better reflect the applied doses of particulate matter 2.5 in the in vitro model. This includes further explanation and discussion of the rationale behind dosage application in vitro, together with further considerations and limitations with physiological translation of this in vitro study to real-life scenarios. This includes; title, methods and discussion.

